# Time-resolved growth of diverse human-associated Akkermansia on human milk oligosaccharides

**DOI:** 10.1101/2025.07.07.663590

**Authors:** Ashwana D. Fricker, Abigail L. Barnes, Arno Henzi, Luis E. Duran, Gilberto E. Flores

## Abstract

The infant gut microbiota is strongly influenced by human milk oligosaccharides (HMOs), a set of glycans that comprise a large constituent of milk and reach the large intestine intact. During growth on HMOs, bacteria produce beneficial metabolites including short chain fatty acids (SCFAs) that are important for host health. Select gut microorganisms have unique sets of enzymes capable of catabolizing distinct HMOs leading to host-specific differences in glycan access, and ultimately differences in SCFA production. Here we cultivated three species of human-associated *Akkermansia,* an early life commensal that is correlated with a healthy metabolic status in adults, on five individual HMOs in two different media backgrounds. Analysis of growth rates, growth yield, metabolic output, and individual HMO consumption through time revealed differences across species that was influenced by growth media. Most notably, *A. biwaensis* CSUN-19 has robust growth in both media backgrounds paired with nearly complete degradation of all HMOs. Across all conditions, overall SCFA production was generally commensurate with growth, but most strikingly, A. *muciniphila* MucT and A. *biwaensis* CSUN-19 produced succinate only when grown in the presence of N-acetyl glucosamine, but not with mucin. The third organism tested, *A. massiliensis* CSUN-17 had weaker growth, lower degradation of HMOs, but higher production of propionate in media containing N-acetyl glucosamine. Interactions between *Akkermansia* and HMOs can influence colonization of other early life commensals, potentially influencing health outcomes throughout life. This study highlights the importance of characterizing growth of individual *Akkermansia* species on distinct HMO leading to fermentation into organic acids.

**IMPORTANCE:** *Akkermansia* are a widely distributed bacterial genus found in the healthy human gut that are capable of degrading host-produced glycans including human milk oligosaccharides (HMOs). Previous end-point experiments demonstrated varying degradation efficiencies across *Akkermansia* species with *A.biwaensis* displaying enhanced growth on multiple HMOs. However, the temporal dynamics and growth preferences when offered substrate choice across the lineage are unknown. Here, we characterized the temporal growth dynamics, HMO catabolism, and metabolic output of three *Akkermansia* species across five HMOs and two media backgrounds. Specifically, we demonstrate that one species, *A. biwaensis* CSUN-19, has robust growth independent of media background with nearly complete degradation of all HMOs tested. Overall, the species-, HMO-, and media-specific response of *Akkermansia* may impact the colonization success of each species, ultimately influencing host-microbe and microbe-microbe interactions in the developing infant gut microbiome.

## INTRODUCTION

Human milk is a unique and highly complex fluid that provides all the necessary nutritional requirements for growth of a developing infant (Smilowitz *et al*. 2014; Kirmiz *et al*. 2018). In addition to nutrition, milk provides important bioactive components including sugars, lipids, enzymes, immunoglobulins, hormones, vitamins, and minerals that provide a range of benefits to the growing infant (Smilowitz *et al*. 2014). Sugars in human milk are highly abundant and can be found as lactose, free human milk oligosaccharides (HMOs), or conjugated to proteins or lipids (Kirmiz *et al*. 2018). HMOs are the third most abundant component of human milk (5-15 g/L) following lactose and lipids (Petherick 2010; Bode 2012). HMOs are composed of a pool of over 200 intricate individual sugars, but only five monosaccharides make up all HMO structures. These are glucose, galactose, N-acetylglucosamine (GlcNAc), fucose, and N-acetylneuraminic acid (NeuAc, sialic acid). Structurally, HMOs consist of a lactose core with the aforementioned monomers linked to this core, resulting in structures that are linear or branched and can be complex or relatively simple (Wu *et al*. 2010). Simple yet abundant HMOs include 2’-fucosyllactose (2’-FL), 3-fucosyllactose (3-FL), lacto-N-tetraose (LNT), lacto-N-neotetraose (LNnT), and 6’-sialyllactose (6’-SL) (Bode 2012; Kirmiz *et al*. 2018). Interestingly, HMOs are not a nutrient source for the developing infant, instead these plentiful sugars influence the assembly of the infant gut microbiome by inhibiting pathogen colonization and enriching for beneficial bacteria (De Leoz *et al*. 2015; Lewis *et al*. 2015; Pacheco *et al*. 2015; Davis *et al*. 2016; Moossavi *et al*. 2018).

The infant microbiome is typically dominated by HMO catabolizing bacteria, most notably from the *Bifidobacterium* genus (reviewed in (Milani *et al*. 2017)). While *Bifidobacterium* are typically dominant in these communities, other bacteria with HMO degrading capabilities are also commonly present (Milani *et al*. 2017). Another genus of organisms present and capable of growing in early life is *Akkermansia*, a lineage of mucin-degrading specialists that is largely regarded as beneficial bacteria in adults (Derrien *et al*. 2004; Cani *et al*. 2022). All known species of *Akkermansia* have a large repertoire of glycoside hydrolase (GHs) enzymes that are involved in mucin catabolism. However, given the structural and compositional similarity between mucin and HMOs (Podolsky 1985; Bode 2012), GHs implicated in mucin catabolism in *Akkermansia* are also likely capable of breaking down HMOs. Importantly, individual species of *Akkermansia* have different sets of GHs, suggesting differences in catabolic capacity across species (Luna *et al*. 2022; Padilla *et al*. 2024). While the original description of growth on HMOs used *A. muciniphila* (Kostopoulos *et al*. 2020), our lab has recently demonstrated end-point growth differences across *Akkermansia* species when grown on individual HMOs and mucin (Luna *et al*. 2022), likely due to the presence and regulation of individual enzymes (Padilla *et al*. 2024).

Given the end-point differences in growth and HMO consumption across *Akkermansia* species in a mucin background, we wanted to examine the temporal dynamics of HMO degradation in different media backgrounds. We hypothesized that each species would consume HMOs at different rates and produce different concentrations of short chain fatty acids (SCFA) as fermentation products. We first determined growth of representative isolates from each of the three named human-associated *Akkermansia* species on five individual HMOs. Subsequently, we selected six time points throughout growth for SCFA and catabolite quantification to identify when individual sugars are digested. Further, to disentangle the response on HMOs from the other sugars present in mucin, synthetic media containing GlcNAc instead of mucin was tested in parallel. From this, we demonstrate differences in growth on individual HMOs, of which one species, *A. biwaensis*, has a robust growth response on multiple HMOs. The ability of *A. biwaensis* to robustly grow on multiple HMO across two media demonstrate its importance for further study as a potential probiotic for infants.

## MATERIALS AND METHODS

### Growth on individual HMOs

*Akkermansia sp.* were grown on basal tryptone threonine medium (BTTM, based on (Ottman *et al*. 2017)) in the presence of 10mM GlcNAc or 0.5% mucin and HMOs, conditions for which we have previously demonstrated robust growth (Ottman *et al*. 2017; Luna *et al*. 2022; Padilla *et al*. 2024). Prior to inoculation, cultures were subcultured twice from glycerol freezer stocks at a 10% and 2% inoculum, respectively, in 5 mL BTTM with 0.4% v/v soluble type III porcine gastric mucin (Sigma-Aldrich) prepared as described previously with overnight growth at 37°C (Luna *et al*. 2022; Fricker *et al*. 2024). Prior to seeding into the experiment, each species was normalized to OD_600nm_ 0.5 with carbon-free basal medium and subsequently inoculated 1:10 into BTTM containing the respective carbon sources. Each tube of BTTM contained a final concentration of 4mM HMOs (2’-FL, 3-FL, 6’-SL, LNT, LNnT) or lactose and 0.4% mucin or 8mM GlcNAc (Supp Table 1). During media preparation, HMOs, lactose, milliQ water, mucin, and GlcNAc were filter sterilized (UNIFLO^TM^ 13mm 0.2μM PES Filter Media, Whatman^TM^). All cultures were prepared and maintained in a Vinyl Anaerobic Chamber (Coy Laboratory Products, Incorporated, Grass Lake, MI) with a headspace of 80% N_2_, 15% CO_2_, and 5% H_2_.

To monitor growth, OD_600nm_ was read hourly using a SpectroSTAR nano spectrophotometer (BMG Labtech, Germany) in clear bottom 96-well plates (Falcon, USA) containing 200 μL of the respective cultures. After inoculation, plates were sealed with a Breathe-easy film ® (Diversified Biotech) to allow for gas exchange, and plates were maintained static at 37°C except for a 10 second orbital shake prior to each hourly read. All experimental cultures were grown in triplicate and experiments were repeated four times.

For experiments involving metabolite detection and tracking of HMO degradation, after preparing cultures, 0.5 mL aliquots were distributed into 2mL microfuge tubes and maintained static separately at 37°C, anaerobically. At the indicated time points, each aliquot was centrifuged at 15,000 rcf for 5 minutes and supernatants were stored at –20°C until further processing (below).

Doubling time and hour at maximum slope was calculated using a rolling slope method. Slopes were calculated across every 5-hour window and the maximum slope during exponential growth was used to determine the doubling time by calculating the ln(2)/slope using the *zoo* package (v.1.8-12) in R. Hypothesis testing was performed using Kruskal-Wallis followed by a post-hoc Dunn’s test using the *rstatix* package (0.7.2). R scripts are provided as Supp Files 1-3. *Metabolite and HMO profiling*

To determine levels of SCFAs (acetate, propionate, and succinate), sugars (lactose, GlcNAc, fucose, sialic acid), and HMOs (2’-FL, 3-FL, 6’-SL, LNT, LNnT) samples were analyzed using high-performance liquid chromatography (HPLC). Prior to analysis, samples maintained at –20°C were filtered (Millex®, 13mm, 0.2uM PTFE Membrane, Millipore) to prevent clogging of the column. Filtered supernatants were measured using an Agilent 1260 HPLC system (Agilent, USA) equipped with a Varian Metacarb 67H column (300 by 6.5 mm) maintained at 45°C with 5 mM sulfuric acid at a flow rate of a 0.8ml/min. A refractive index detector at 35°C was used to measure glycans and metabolites. Concentrations were determined by comparing the area under the curve to six-point standards for each compound of interest; a chemical list of all standards is provided in Supp Table 2. Graphs were generated using the *ggplot2* package (v.3.5.1) and significant differences between samples were calculated using the *rstatix* package in R v.4.2.3.

### Vitamin B12 Bioassay

To determine if vitamin B12 was present in the mucin used as a growth substrate, we used the *Lactobacillus leichmannii* ATCC 7830 bioassay. For this bioassay, the original protocol was modified and described in previous work (Valu 1965; Kirmiz *et al*. 2020). Briefly, sterile vitamin B12 assay medium (BD Difco) with standard concentrations (0 to 2.5 ng/ml) of cyanocobalamin (Sigma-Aldrich) or different batches of mucin to a final concentration of 0.05% (note: this represents a 10x lower concentration than in media used to grow *Akkermansia*). Cyanocobalamin and mucin were filter sterilized prior to addition to cooled assay media (UNIFLO^TM^ 13mm 0.2μM PES Filter Media, Whatman^TM^). After media preparation, overnight cultures of *L. leichmannii* grown in MRS at 37°C in 5% CO_2_ were pelleted at 15,000 rcf for 3 minutes and washed 3X with sterile Milli-Q water. Pellets were resuspended in sterile water and incubated at 4°C for one hour. This culture was diluted 1:10 with sterile water before inoculating 1:100 into 1mL vitamin B12 assay medium (BD Difco) in a 48-well plate. Cultures were incubated for 20 hours at 37°C and 5% CO_2_. Growth was measured with a Spectromax M5 (Molecular Devices, USA) at OD_600nm_. Values from the Spectromax plate reader were converted to true OD_600nm_ using a linear conversion.

## RESULTS

### 3.1 Growth on HMO is species and media dependent

To identify differences in growth patterns on HMOs across species of human-associated *Akkermansia*, we first cultivated a representative of each named species on HMOs in the presence of mucin. A time-resolved approach revealed differences in species-level growth that was HMO-dependent (Figure 1).

**Figure 1.**
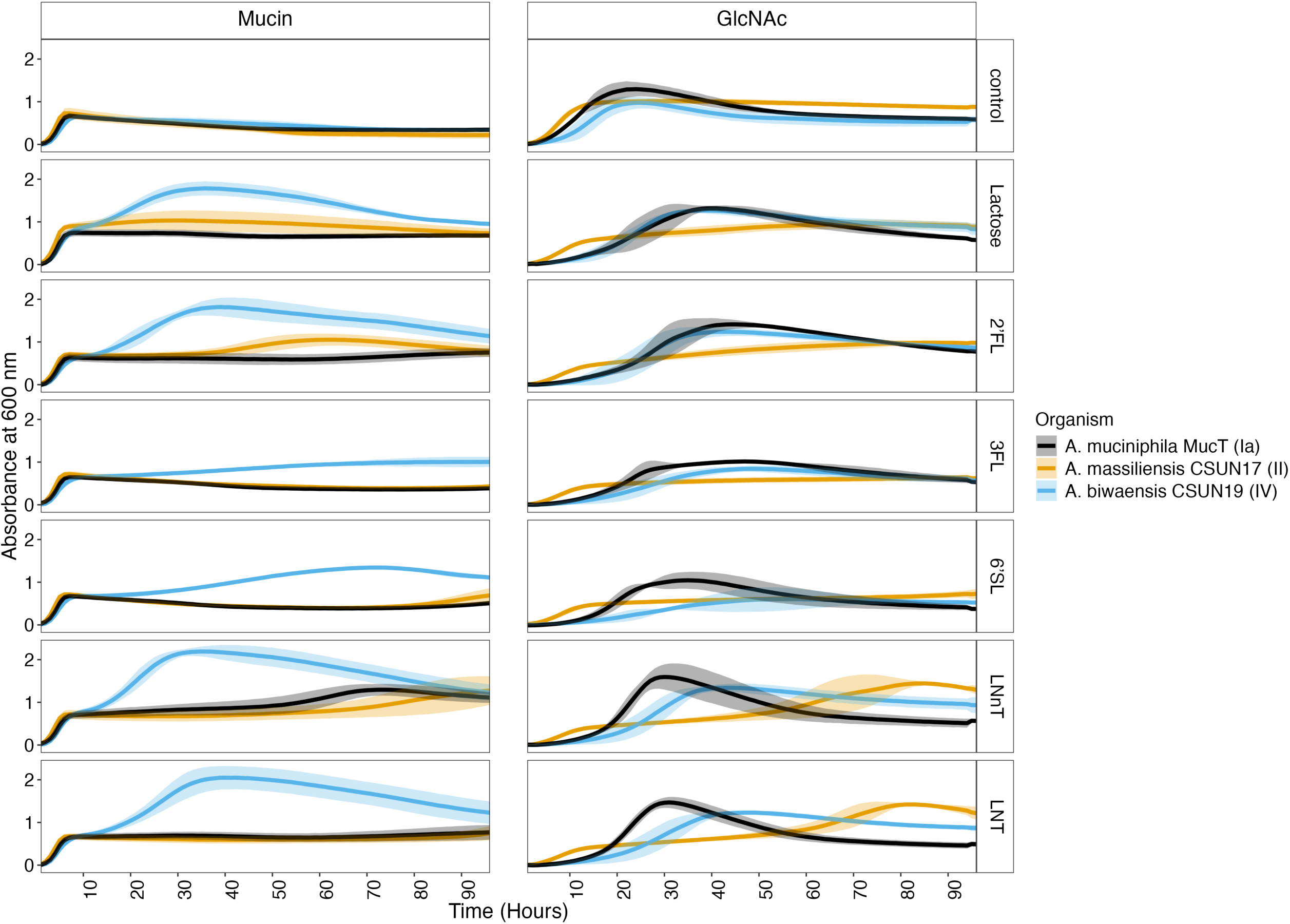
*Akkermansia* species demonstrate different growth responses to individual HMOs through time in a media-dependent manner. OD600 was measured hourly for 96h for three species across five HMOs in the presence of either mucin or N-acetylglucosamine (GlcNAc). To normalize OD, the initial time point (t = 0h) was subtracted from all subsequent reads. Values represent the average of three to four biological replicates, each performed in triplicate. Error bars represent standard deviation as calculated in R.

#### 3.1.1 Species vs Species

In the mucin medium lacking HMOs, all three representatives had equal growth, reaching equal OD_600_ maxima within 6 hours with a slight delay in *A. biwaensis* CSUN-19 (Supp Figure 1A & E). However, across all five HMOs tested in a mucin background, *A. biwaensis* CSUN-19 had the most robust growth, with significantly higher OD_600_ maxima compared to the other species tested (p<0.01, Supp Figure 1A & D).

To eliminate the complexity of mucin potentially influencing growth, the same experiment was repeated in the absence of mucin. Instead, GlcNAc was included in the medium to alleviate the auxotrophy for this substrate. As a control, glucose was provided as a carbon and energy source, which equally supported growth of all species (Figure 1B, Supp Figure 1A & D). In the presence of GlcNAc, the influence of the HMOs varied drastically across species (Figure 1B). For example, both *A. muciniphila* Muc^T^ and *A. biwaensis* CSUN-19 had similar growth patterns on lactose, 2’-FL, and 3-FL, whereas *A. massiliensis* CSUN-17 seemed to have an initial early growth phase with a continued gradual increase in OD_600_ over the remaining time points on these same three sugars. Despite the early and continued growth on these three sugars, *A. massiliensis* CSUN-17 had the lowest OD_600_ maximum (Supp Figure 1A). Comparatively, each species had different growth patterns on 6’-SL; on this HMO in the presence of GlcNAc, *A. muciniphila* Muc^T^ grew abundantly after a ∼12h lag and had the highest maximum OD_600_ whereas both *A. massiliensis* CSUN-17 and *A. biwaensis* CSUN-19 had gradual increases in OD_600_. Across both LNT and LNnT, the patterns of growth appeared similar, with *A. massiliensis* CSUN-17 demonstrating biphasic growth, and *A. muciniphila* Muc^T^ and *A. biwanesis* CSUN-19 growing relatively quickly after a long lag. Notably, all organisms had the same final growth yield on LNnT and similar growth on LNT in a GlcNAc, but not mucin, background (Supp Figure 1A).

All isolates had a slight decrease in OD_600_ during late stages of growth on mucin alone, which did not occur in the presence of HMOs or lactose. This led to significant differences in the growth with additional sugars compared to mucin alone at both 24 and 48h (p<0.05, Supp Figure 2), where the general trends are in line with previous work (Luna *et al*. 2022). Curiously, during growth on GlcNAc across multiple HMOs, *A. muciniphila* Muc^T^ had a marked decrease in OD_600_ after peak growth, although this effect was most pronounced in media with LNT or LNnT (Figure 1).

**Figure 2.**
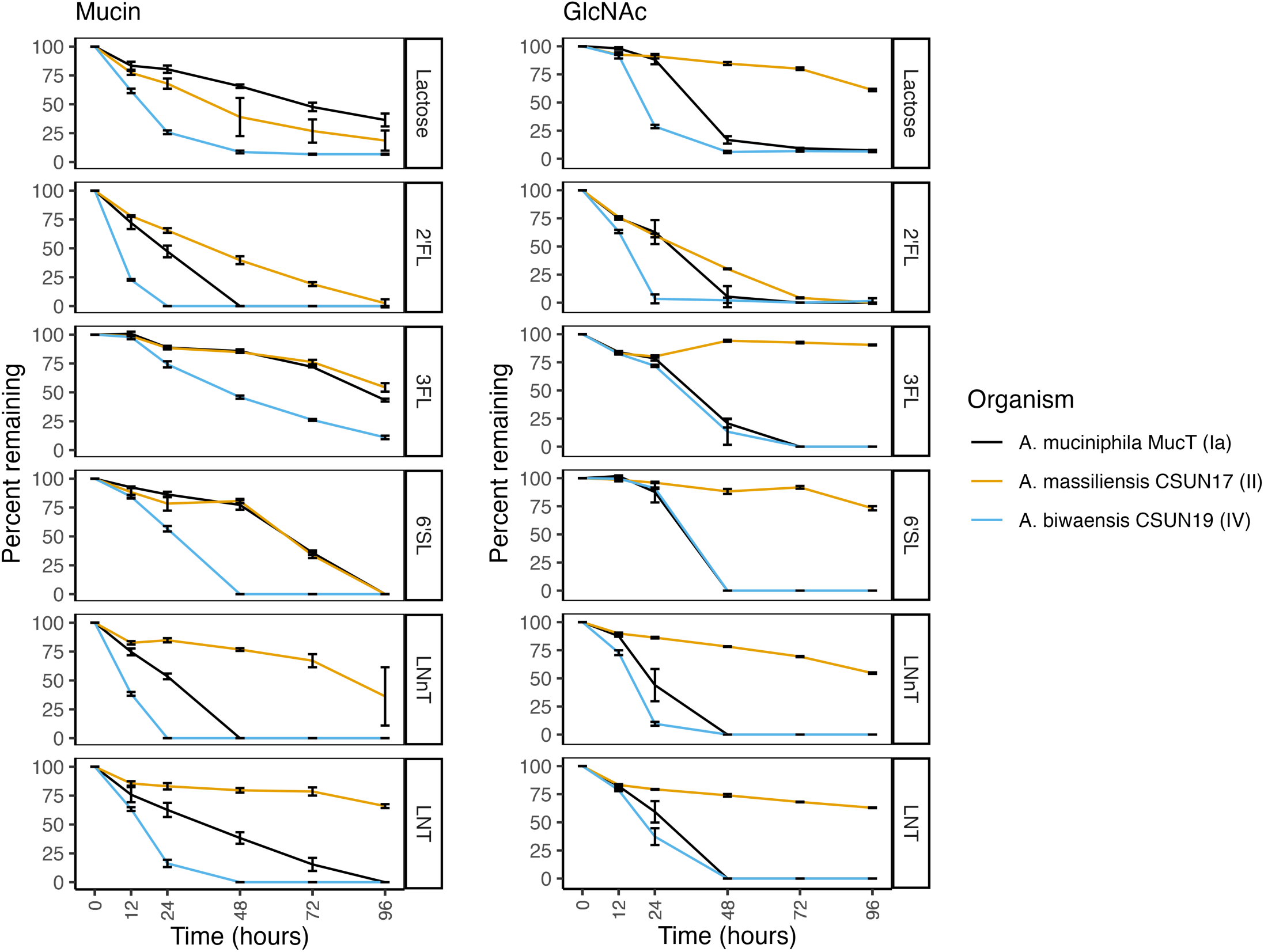
HMO degradation rates by *Akkermansia* are species– and media-dependent. Percent decrease as compared to time zero of each HMO was determined by HPLC and captured at 5 time points during growth on either mucin (left) or GlcNAc (right). Error bars represent standard deviation of three technical replicates.

#### 3.1.2 Media v Media

The influence of the media was different across species; for example, *A. muciniphila* Muc^T^ had generally better growth on GlcNAc, *A. massiliensis* CSUN-17 had approximately equal growth on both media, and *A. biwaesnsis* CSUN-19 grew better overall in a mucin background (Figure 1). This trend was most noticeable when comparing the maximum OD_600_ and the time at which this maximum was reached (Supp Figure 1B & E).

#### 3.1.3 HMO v HMO

Within each species, the ability to grow on different HMOs was compared. In the mucin background, *A. massiliensis* CSUN-17 had similar OD_600_ maxima when presented with different HMOs whereas in a GlcNAc background, there was much greater variability across HMOs (Supp Figure 1C & F). All three species had higher OD_600_ maxima on LNnT and LNT as compared to the HMO that supported the least amount of growth, 3-FL (Supp Figure 1C).

To quantify diauxic growth patterns, the maximum doubling time and the midpoint hour at which the slope was calculated were compared for the first and second log phases, when present. Strikingly, across all species and HMOs in a mucin background, the doubling time during the first log phase was similar (Supp Figure 3). However, there were large differences in the doubling time during the second log phase that was HMO-dependent. This suggests early growth on mucin, and subsequent growth on HMO. However, in a GlcNAc background, only *A. massiliensis* CSUN-17 had diauxic growth or continued increases in growth across multiple HMOs occurring after 48h of incubation (Figure 1). These doubling times for CSUN-17 suggest early growth on GlcNAc and subsequent use of HMO, and points to potential preference for LNT and LNnT.

**Figure 3.**
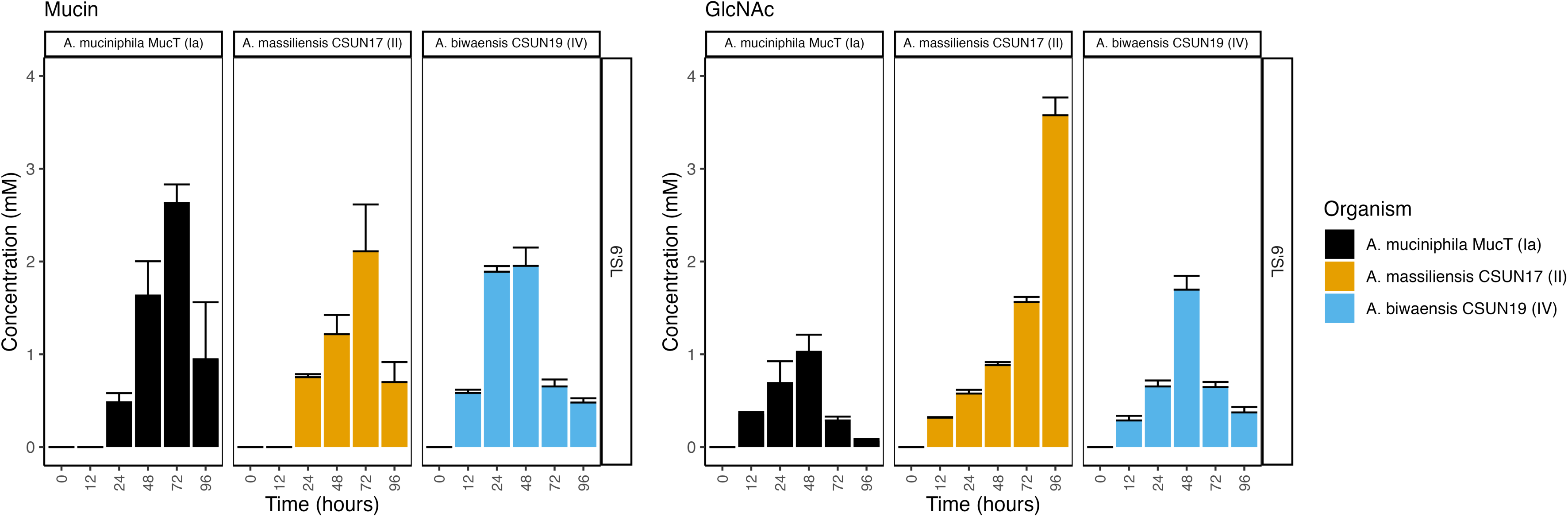
Free sialic acid concentrations peak at 48-72 hours followed by a decrease as Akkermansia deconstruct 6’-siallylactose. Concentrations of sialic acid were determined by HPLC and captured at 6 time points during growth on either mucin (left) or GlcNAc (right). Error bars represent standard deviation of three technical replicates.

### 3.2 Rates of HMO degradation are species and HMO dependent

To determine whether rates of HMO degradation matched growth patterns, culture supernatants were analyzed via HPLC before incubation, at 12h, and subsequently every 24h throughout growth. In a mucin background, the organism that had the most growth across all HMOs, *A. biwaensis* CSUN-19, also degraded each of the HMOs faster than the other species (Figure 2). For example, 2’-FL and LNnT were completely digested by 24h. Similarly, *A. muciniphila*, Muc^T^, catabolized all the 2’-FL and LNnT within 48h. Surprisingly, in the mucin background, all three organisms catabolized 6’-SL, despite weak growth on this HMO by both *A. muciniphila* Muc^T^ and *A. massiliensis* CSUN-17.

When grown with GlcNAc, the patterns of decreasing HMO concentrations were different than during growth on mucin. For example, both *A. muciniphila* Muc^T^ and *A. biwaensis* CSUN-19 had similar patterns of 3-FL and 6’-SL on GlcNAc media, whereas in a mucin background, *A. muciniphila* Muc^T^ more similarly resembled *A. massiliensis* CSUN-17 on these HMOs. Across all HMOs except 2’-FL, *A. massiliensis* CSUN-17 did not appear to robustly degrade any in the absence of mucin, suggesting this organism may have primarily used GlcNAc as a carbon and energy source. To address this, levels of GlcNAc were measured via HPLC (Supp Figure 4). There was a large drop in the levels of GlcNAc within the first 12h, whereas *A. muciniphila* Muc^T^ and *A. biwaensis* CSUN-19 appeared to use GlcNAC within the 12-24h window, suggesting a need to acclimate to the absence of mucin. In addition, while all the GlcNAc was consumed by both *A. muciniphila* Muc^T^ and *A. biwaensis* CSUN-19 within 48h across all HMOs (with the singular exception of *A. biwaensis* CSUN-19 grown on LNT), there was residual GlcNAc in all *A. massiliensis* CSUN-17 cultures by the end of the experiment (96h).

**Figure 4.**
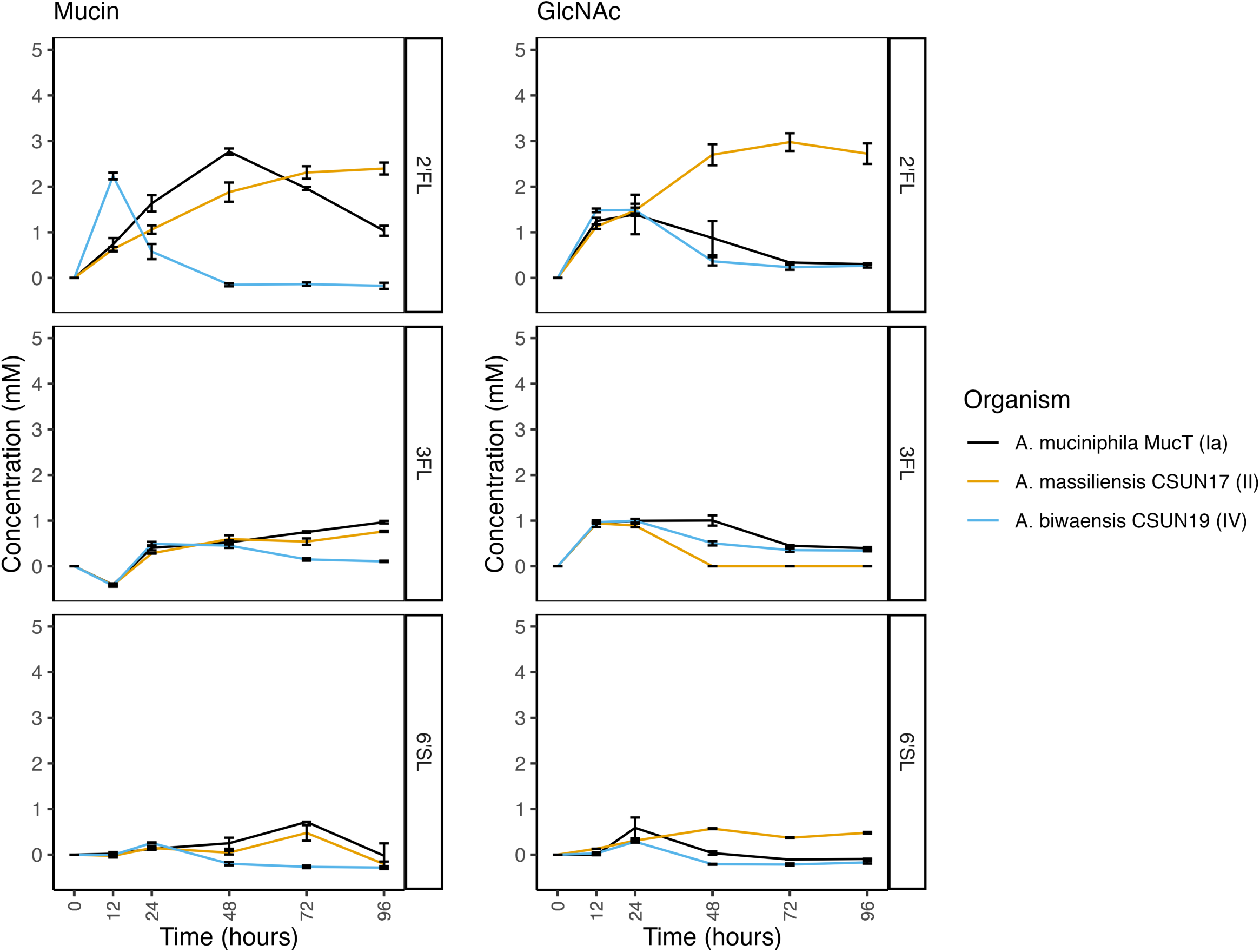
Increase in lactose is species-dependent. Concentrations of lactose were determined by HPLC and captured at 6 time points during growth on either mucin (left) or GlcNAc (right). Error bars represent standard deviation of three technical replicates.

#### Byproducts of HMO breakdown suggest metabolism of intermediates

In hand with breakdown of HMOs, levels of sugar breakdown products were determined via HPLC. Trace amounts of fucose (less than 1 mM) were detected in most cultures grown on fucosyllated HMOs, however, the levels were close to the detection limit (data not shown). Perhaps surprisingly, although the concentration of NeuAc increased over the first 48h-72h in all three organisms across both media backgrounds supplemented with 6’-SL, the concentration decreased at later growth stages with the exception of *A. massiliensis* CSUN-17 in a GlcNAc background (Figure 3).

Given that lactose forms the backbone of all HMOs, we also measured free lactose in the culture medium for all three species across both media backgrounds (Figure 4). However, lactose peaks overlapped with unidentified di– or tri-saccharide peaks in media containing LNT and LNnT, precluding lactose measurements for those HMOs. Lactose was transiently detected in nearly all cultures of *A. biwaensis* CSUN-19, likely due to quick consumption. In cultures of *A. muciniphila* Muc^T^ grown on 2’-FL, levels of lactose peaked at 48h in a mucin background and at 24h in a GlcNAc background, and then slowly diminished throughout the rest of the experiment. For this organism on 3-FL and 6’-SL, lactose concentrations never accrued to more than 1mM. All cultures of *A. massiliensis* CSUN-17 appeared to let lactose accumulate in the culture medium throughout the course of the experiment during growth on 2’-FL in both media backgrounds.

#### A. biwaensis produces greater amounts of SCFAs

To identify whether individual HMOs led to differences in metabolic output, SCFAs were measured from the same six time points (0, 12, 24, 48, 72, and 96 hours) of each isolate. For all three organisms under mucin only conditions, approximately 5 mM acetate and 3mM propionate were produced, levels which fluctuated marginally after 12h (Figure 5). Similarly, under growth with lactose and mucin, both *A. massiliensis* CSUN-17 and *A. biwaensis* CSUN-19 produced similar levels of acetate and propionate, whereas *A. muciniphila* Muc^T^ produced significantly less propionate (p<0.05 at 96h). Across all HMOs in a mucin background, *A. biwaensis* CSUN-19 produced the most SCFAs. Notably, despite ample production of acetate and propionate during growth on lactose, *A. massiliensis* CSUN-17 demonstrated equal or weaker production of these SCFAs as compared to *A. muciniphila* Muc^T^ when grown with HMO. Interestingly, both *A. biwaensis* CSUN-19 and *A. muciniphila* Muc^T^ had small amounts of succinate production which built up slightly in the culture medium before disappearing.

**Figure 5.**
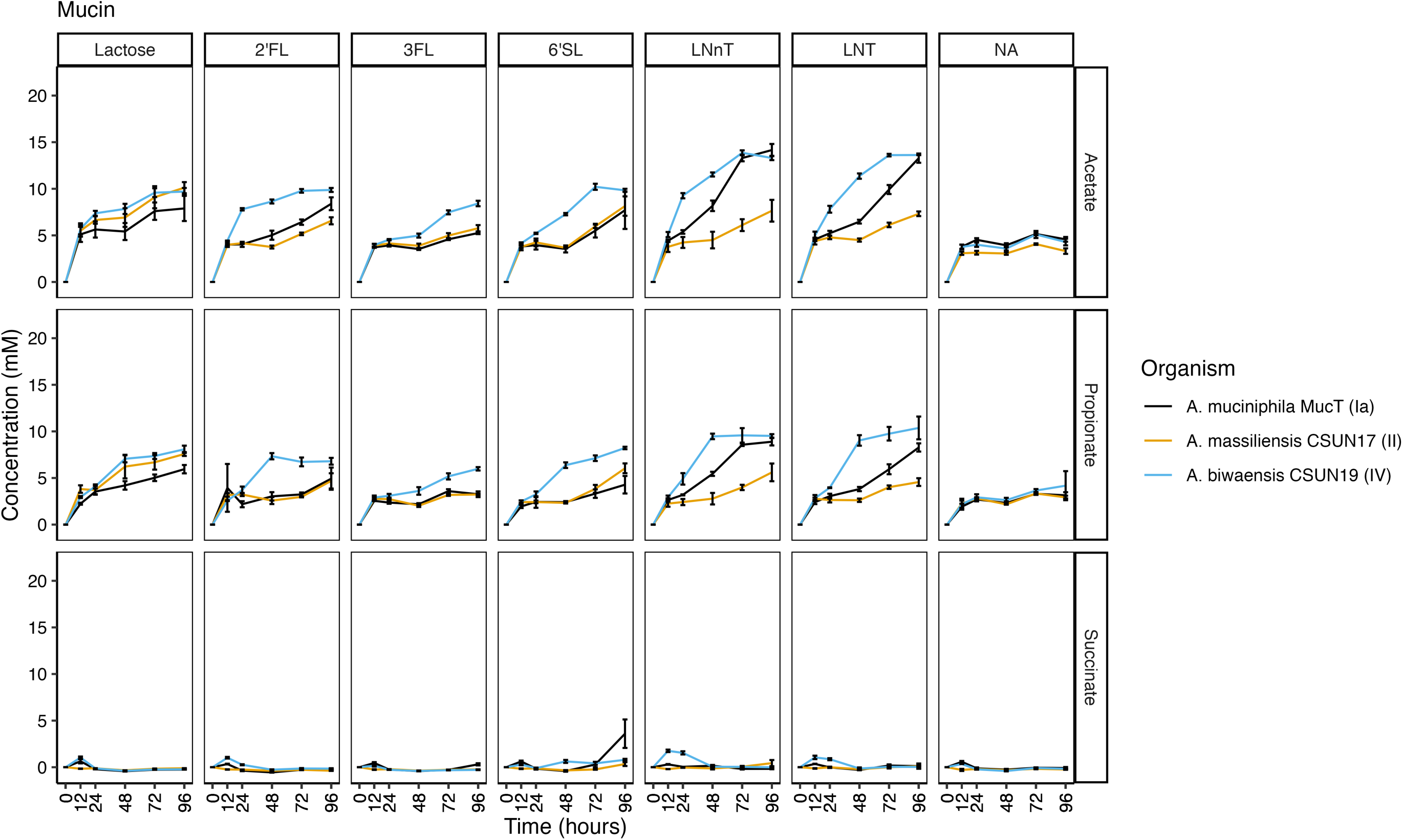
Short chain fatty acids are synthesized continuously throughout growth on mucin and HMOs. Concentration of acetate, succinate, and propionate during growth on mucin was determined by HPLC and captured at 6 time points. Error bars represent standard deviation of three technical replicates.

The same SCFAs were detected in cultures grown with a GlcNAc background. Similar to growth on mucin alone, all three organisms had the same amount of acetate accumulate, reaching around 10mM in the glucose + GlcNAc condition. However, *A. massiliensis* CSUN-17 produced higher levels of propionate and trace levels of succinate under these conditions (Figure 6). A similar pattern emerged during growth on lactose and all HMOs in a GlcNAc background in which *A. massiliensis* CSUN-17 had significantly higher levels of propionate, but lower levels of succinate as compared to *A. muciniphila* Muc^T^ and *A. biwaensis* CSUN-19 (p<0.05 at 96h). Comparing the two high-acetate producers, *A. biwaensis* CSUN-19 and *A. muciniphila* Muc^T^, revealed differences during growth on both LNT and LNnT, where despite equal production of acetate and propionate, *A. biwaensis* CSUN-19 had higher levels of succinate produced.

**Figure 6.**
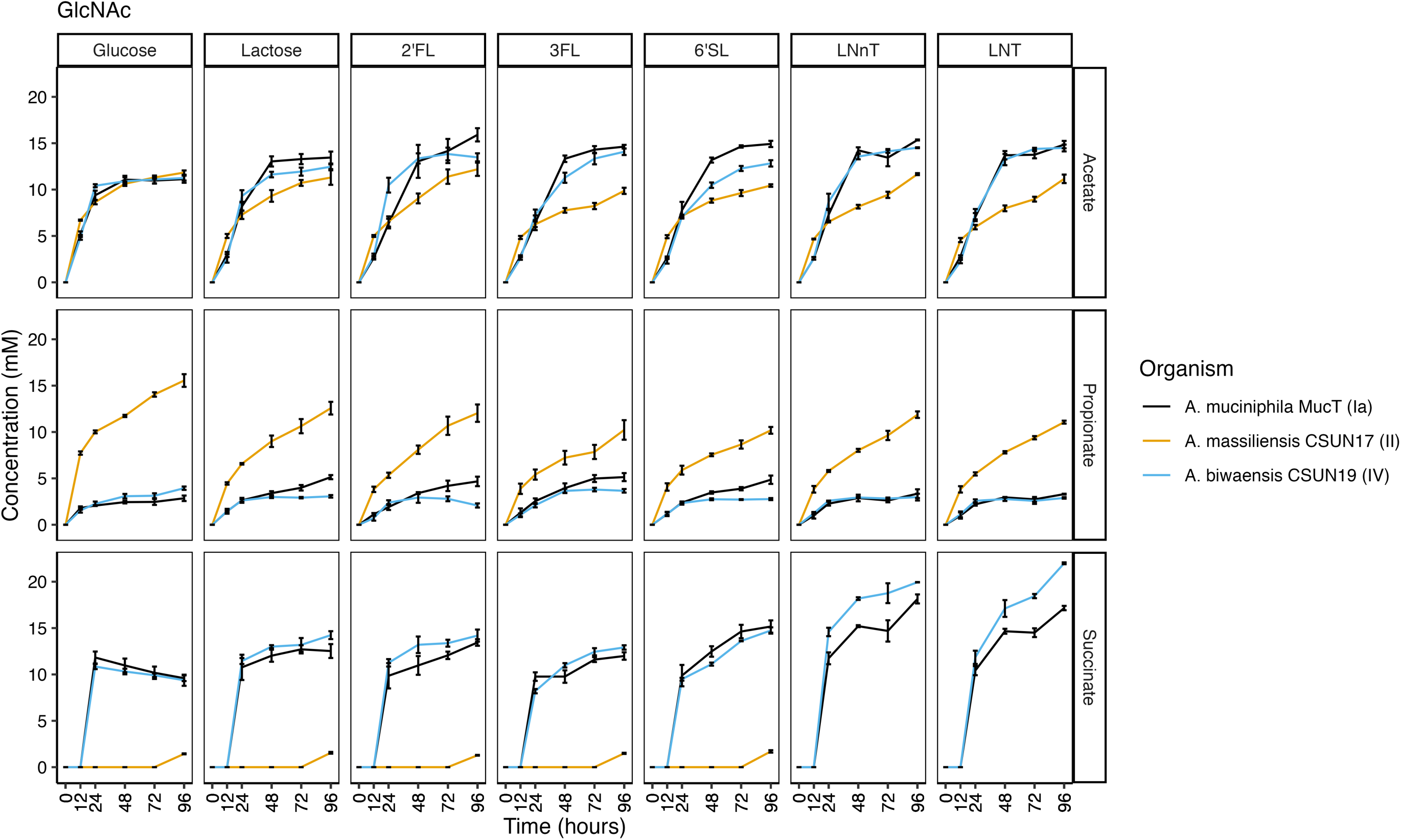
Short chain fatty acids are synthesized continuously throughout growth growth on GlcNAc and HMOs. Concentration of acetate, succinate, and propionate during growth on GlcNAc was determined by HPLC and captured at 6 time points. Error bars represent standard deviation of three technical replicates.

The increase and then subsequent decrease of succinate that paralleled increasing propionate concentrations in *A. muciniphila* Muc^T^ and *A. biwaensis* CSUN-19 in the mucin but not the GlcNAc media suggested potential vitamin B12 scavenging from mucin (Kirmiz *et al*. 2020; Mok *et al*. 2020). To determine if vitamin B12 was present in mucin, we used a *Lactobacillus leichmannii* ATCC 7830 bioassay to verify presence of this vitamin. Mucin supported the growth of the vitamin B12 auxotroph, thus confirming the presence of vitamin B12 in mucin preparations (Supp Figure 5).

## DISCUSSION

Human milk oligosaccharides have intricate interactions with early life colonizers of the human gut microbiota, such as members of the genera *Bifidobacterium*, *Akkermansia*, and *Bacteroides*. Our prior work demonstrated that individual *Akkermansia* species have unique responses to growth across multiple HMOs, partially mediated by differences in the suite of GH enzymes encoded in their genomes (Luna *et al*. 2022; Padilla *et al*. 2024). In turn, these differences likely affect the ecology of *Akkermansia* in a species dependent manner. Here, we extend our initial screening of *Akkermansia* growth on HMOs by analyzing the temporal growth dynamics, HMO degradation profiles, and metabolic output across two media backgrounds. We found that *A. biwaensis* CSUN-19 has robust growth in a mucin-media background that is paired with quick degradation of the corresponding HMOs and high production of health-promoting SCFAs. Comparatively, in this same background, the growth of *A. mucinphila* Muc^T^ and *A. massiliensis* CSUN-17 was dependent on the provided HMO, with overall moderate HMO degradation and SCFA production. Growth dynamics changed drastically in a GlcNAc background, wherein *A. muciniphila* Muc^T^ had growth, HMO degradation, and SCFA production patterns similar to *A. biwaensis* CSUN-19. Although *A. massiliensis* CSUN-17 had early growth in this background, the minimal degradation of the HMOs and lower SCFA production suggests overall weak ability for HMO usage under these conditions. Overall, these findings demonstrate that the response to HMOs is species-, HMO-, and media-specific, which may impact the colonization success and overall ecology of *Akkermansia* in the developing infant gut.

Previously, we established HMO degradative capabilities of diverse human associated *Akkermansia* using 48-hour end-point experiments in a mucin medium background (Luna *et al*. 2022). Here, we extend those findings to include time-resolved growth dynamics and quantitative metabolite production of three *Akkermansia* species in two media backgrounds. While there are subtle differences in the results between our two experiments (most notably, *A. biwaensis* CSUN-19 consumed more HMOs after 48 hours in this experiment than in our previous study), we continue to observe that *A. biwaensis* CSUN-19 has the greatest growth rates and yields when grown on HMOs. While *A. biwaensis* does not appear to be the dominant species of *Akkermansia* in adult human populations (Mueller *et al*. 2024), it has been isolated from adults across the globe (Becken *et al*. 2021; Luna *et al*. 2022; Kobayashi *et al*. 2023). Additionally, the original discovery of this *Akkermansia* lineage (formally known as phylogroup AmIV) was made from metagenomic sequences originating from a population of children aged 2-9 years living in southern California, USA (Herman *et al*. 2020; Kirmiz *et al*. 2020). Given the rapid growth and deconstruction of several core HMO structures, future studies should more directly address the prevalence and distribution of *A. biwaensis* in diverse infant populations.

*Akkermansia* are mucin-degrading specialists in part because of their inability to aminate fructose-6-phosphate (Fru6P) to form glucosamine-6-phosphate (GlcN6P), a precursor to N-acetyl-D-glucosamine (GlcNAc) (McCown, Winkler and Breaker 2012). GlcNAc is needed for peptidoglycan biosynthesis (Garde, Chodisetti and Reddy 2021). Fortunately, mucin glycoproteins are rich in GlcNAc and provide a reliable source for this essential nutrient *in vivo* (Tailford *et al*. 2015). In the laboratory, this auxotrophy can be overcome by providing mucin or a sugar amine (i.e., GlcNAc or GalNAc) directly in media formulations. Here, in line with previously reported differences across *Akkermansia* species during growth on synthetic versus mucin media (Becken *et al*. 2021), we also observe species-specific effects of GlcNAc on growth. For example, *A. massiliensis* CSUN-17 had very similar first-log phase responses to growth in all conditions tested and the largest drop in GlcNAc concentrations within the first twelve hours, suggesting the initial growth in the GlcNAc background is using this sugar amine alone rather than the supplemented sugars or HMOs. Ability to use GlcNAc may be dependent on transport mechanisms for this sugar amine: genes that code for GlcNAc transport could not be identified for *A. muciniphila*, suggesting reliance on other sugar transport mechanisms (van der Ark *et al*. 2018). As GlcNAc supplemented media is suitable for industrial scale production of *A. muciniphila* (Plovier *et al*. 2017; van der Ark *et al*. 2018), understanding how sugar amines differentially support growth may be important for development of other *Akkermansia* species.

HMO degradation by bacteria is often initiated by removal of the terminal sugar on the non-reducing end of the polysaccharide by exo-acting GH enzymes (Shuoker *et al*. 2023). In the case of the trisaccharide HMOs tested here, this initial hydrolysis would result in free fucose (2’-FL and 3-FL) or sialic acid (6’-SL) and lactose. Indeed, based on detectable levels of fucose, sialic acid, and lactose in spent culture media through time, coupled with the previous characterization of several fucosidase and sialidase enzymes with these activities (Shuoker *et al*. 2023; Bakshani *et al*. 2025), it appears that *Akkermansia* utilize this sequential degradation strategy for these HMOs. However, while degradation of tetrasaccharide HMOs like LNT and LNnT can also be sequential, non-sequential degradation enabled by endo-acting GH enzymes has also been observed in other early life colonizers (reviewed in (Sakanaka *et al*. 2019)). While mediated by different GH enzymes, sequential degradation of LNT and LNnT would result in the trisaccharide lacto-N-triose II (LNT-II) and Gal (Sakanaka *et al*. 2019). Sequential degradation of LNT and LNnT is common among *Bifidobacterium* species given the for this process (Sakanaka *et al*. 2019). In contrast, non-sequential degradation could result in production of two disaccharides; for LNT these are lacto-N-biose (LNB) and lactose, while for LNnT they would be N-acetyllactosamine (LacNAc) and lactose (Asakuma *et al*. 2011). For example, some species of *Bifidobacterium* use an extracellular endo-acting lacto-N-biosidase (GH20) enzyme to cleave LNT into LNB and lactose (Wada *et al*. 2008). When attempting to quantify lactose in our cultures grown on LNT and LNnT, we noticed a small shift in the retention time (< 30 seconds) near where lactose elutes suggesting the presence of a different sugar, possibly LNT-II, LNB (for LNT), or LacNAc (for LNnT). As we are unable to resolve the identity of this peak, we are unsure of the strategy employed by these *Akkermansia*. It is also worth noting that two recombinant, purified GH16 enzymes from *A. muciniphila* Muc^T^ were shown to remove the terminal glucose from both LNT and LNnT resulting in liberation of different trisaccharides (Crouch *et al*. 2020). While we currently do not know the strategy *Akkermansia* use to deconstruct these core HMO structures, future studies will determine these strategies.

Across all three organisms grown on 6’-SL, free sialic acid was detected in the culture media with concentrations peaking at 48-72 hours with one exception, *A. massiliensis* CSUN-17 grown in a GlcNAc background. The accumulation of sialic acid has previously been noted during growth on natural pools of HMOs and individual HMOs, as well as on heavily sialylated mucins (Kostopoulos *et al*. 2020; Luna *et al*. 2022; Shuoker *et al*. 2023). However, we also observe a decrease in sialic acid in the culture media at later time points for most cultures, suggesting breakdown or use of this compound. In host-associated microorganisms, sialic acids can be degraded for energy conservation (Vimr *et al*. 2004) or used to decorate the cell surface to evade host immune detection (Severi, Hood and Thomas 2007). Despite the absence of a canonical N-acylneuraminate (*nan*) catabolic system, it is possible that *Akkermansia* use a different mechanism to degrade or use sialic acid. Of note, *Akkermansia* possess an N-acetylneuraminate lyase gene (e.g., Amuc_1946) that may convert sialic acid to N-acetylmannosamine (ManNAc) and pyruvate. From this, the ManNAc can be converted to UPD-N-acetylglucosamine by a UDP-N-acetyl-D-glucosamine 2-epimerase encoded by Amuc_1947. The resulting UDP-N-acetyl-D-glucosamine can be shunted towards cell wall biosynthesis (NAM or NAG) (Michal and Schomburg 2013). The pyruvate resulting from the initial conversion could supply minimal energy for growth. However, the buildup of sialic acid in the culture medium and consumption of lactose suggests that much of the energy used for growth on 6’-SL likely derives from lactose. Within the context of the gut microbiome, it is possible that the residual sialic acid can be used to support the growth of other gut bacteria (Shuoker *et al*. 2023).

It is also possible that instead of converting sialic acid to useable energy, *Akkermansia* incorporates sialic acid into its cell envelope (e.g., lipopolysaccharide (LPS) and capsular polysaccharide) helping to avoid host immune detection, in line with other host-associated microorganisms (Severi, Hood and Thomas 2007; Garcia-Vello *et al*. 2024). Future work is needed to determine the fate of sialic acid liberated from host produced glycans by *Akkermansia*.

Unexpectedly, in media containing mucin, all cultures produced large amounts of propionate and no succinate, whereas in the GlcNAc medium, only *A. massiliensis* CSUN-17 produced propionate and no succinate. The conversion of propionate to succinate occurs through the vitamin B12-dependent methylmalonyl-CoA synthase (Belzer *et al*. 2017; Kirmiz *et al*. 2020). Our previous work established that *A. massiliensis* CSUN-17, but neither *A. muciniphila* Muc^T^ nor *A. biwaensis* CSUN-19 synthesize vitamin B12 *de novo* (Kirmiz *et al*. 2020). However, others have shown that all *Akkermansia* known to date possess a cobamide remodeling gene, CbiR, that allows them to remove and replace the lower ligand of diverse forms of vitamin B12 resulting in production of a useable form (Mok *et al*. 2020). Since we do not supplement our media with vitamins, these observations suggest that mucin is providing a form of vitamin B12 accessible to *Akkermansia*. Vitamin B12 scavenging from mucin may explain previous inconsistencies and batch-to-batch variation in methionine production, another dependent function, while supporting *A. muciniphila* Muc^T^ growth (Mok *et al*. 2020). This conversion potentially influences microbe-host interactions, as succinate is involved in immunomodulation (Fernández-Veledo and Vendrell 2019) and propionate is linked to appetite suppression, potentially mediated through stimulating GLP-1 secretion (Byrne *et al*. 2015). The ability for some *Akkermansia* (i.e. *A. massiliensis* CSUN-17) to produce propionate in the absence of vitamin B12 may suggest a role in mitigating the effects of obesity and should be further studied. Further, ratios of SCFA production may lead to cross-feeding dynamics with other members of the gut microbiome. For example, the liberation of sugars from mucin by *A. muciniphila* Muc^T^ supports the growth of *E. hallii,* which produces vitamin B12, allowing *A. muciniphila* Muc^T^ to convert succinate to propionate (Belzer *et al*. 2017). These types of co-culture experiments have revealed an upregulation of genes involved in mucin degradation in *A. muciniphila* during growth with other microorganisms (Chia 2018), suggesting a propensity for *Akkermansia* to act as a primary degrader for the community. Subsequently, the *A. muciniphila* co-cultures reveal increased butyrate production in butyrogenic members of the gut microbiota including *A. caccae*, *R. inulinivorans, F. prausnitzii*, and others (Belzer *et al*. 2017; Chia 2018; Pichler *et al*. 2020; Shuoker *et al*. 2023). However, it is unclear whether these scavengers directly consume mucin sugars or act as secondary fermenters consuming acetate released by *Akkermansia*. Further, it is possible that other secondary fermenters of the gut microbiome engage in metabolic cross-feeding by consuming succinate produced by *A. muciniphila* and *A. biwaensis*, as has been observed for other gut microorganisms (Culp and Goodman 2023). Therefore, future work aimed at disentangling the metabolic interactions of *Akkermansia* species and other gut microorganisms is essential to holistically understand their impact on human health.

## CONCLUSIONS

In conclusion, the ability of *Akkermansia* to fully deconstruct mucin oligosaccharides (Bakshani *et al*. 2025) and a variety of core HMO structures uniquely distinguishes them from other members of the infant gut microbiome. However, like other members of the infant gut microbiome, HMO degradation efficiency and potentially the mechanisms employed, are species and strain dependent. How these differences impact the colonization success and *in vivo* activity of each *Akkermansia* in mixed gut communities remains largely unknown. Ongoing and future studies will more directly assess how differences across species may impact host-microbe and microbe-microbe interactions in the context of the infant gut microbiome.

## Author Contributions

Conceptualization, G.E.F.; methodology, A.D.F.; formal analysis and investigation, A.D.F., A.L.B., A.H., L.E.D.; visualization, A.D.F.; writing-preliminary draft, A.D.F., G.E.F.; writing-revision, A.D.F., G.E.F.; supervision, G.E.F. All authors provided critical feedback and helped shape the research, analysis, and manuscript. All authors have read and agreed to the published version of the manuscript.

## Funding

This work was supported by the National Institute of General Medical Sciences (NIGMS) [grant number SC1GM136546] of the National Institutes of Health. The content is solely the responsibility of the authors and does not necessarily represent the official views of the National Institutes of Health.

## FIGURE LEGENDS

**Supplemental Figure 1.** Differences in growth yield are statistically significant across species, HMOs, and media. Maximum growth yield (A,B, C) and the time at which maximum growth was reached (D, E, F) was compared across species (A, D) across background media (B, E), and across HMOs (C, F). Representative strains used in this study are *A. muciniphila* MucT (Ia), *A. massiliensis* CSUN-17 (II), and *A. biwaensis* CSUN-19 (IV). Error bars represent standard deviation of four biological replicates; statistical significance was determined by Dunn’s test with Bonferroni corrections following Kruskal-wallis in R. Symbol styles: (ns) non significant, (*) 0.05, (**) 0.01, (***) 0.001, (****) <0.001.

**Supplemental Figure 2.** Relative growth on HMOs is different across species. Growth on the mucin-only culture was subtracted from growth on an individual HMO at 24 hours (A) and 48 hours (B) for each species. Representative strains used in this study are *A. muciniphila* MucT (Ia), *A. massiliensis* CSUN-17 (II), and *A. biwaensis* CSUN-19 (IV). Error bars represent standard deviation of four biological replicates; statistical significance was determined by Dunn’s test with Bonferroni corrections following Kruskal-wallis in R. Symbol styles: (ns) non significant, (*) 0.05, (**) 0.01, (***) 0.001, (****) <0.001.

**Supplemental Figure 3.** Despite presence of HMOs, in a mucin background, the doubling time and max slope for the first log phase are the same within a species. Representative strains used in this study are *A. muciniphila* MucT (Ia), *A. massiliensis* CSUN-17 (II), and *A. biwaensis* CSUN-19 (IV). Error bars represent standard deviation of three to four biological replicates; statistical significance was determined by Dunn’s test with Bonferroni corrections following Kruskal-wallis in R. Symbol styles: (ns) non significant, (*) 0.05, (**) 0.01, (***) 0.001, (****) <0.001.

**Supplemental Figure 4.** Decreases in GlcNAc in all strains when grown in this background that is rate-dependent. Concentrations at each of 6 time points was determined by HPLC during growth on GlcNAc. Error bars represent standard deviation of three technical replicates.

**Supplemental Figure 5.** Mucin contains vitamin B12, supporting the growth of *Lactobacillus leichmanii*. Growth yield of *L. leichmanii* after cultivation with increasing concentrations of vitamin B12 or mucin. Error bars represent standard deviation of three technical replicates experiment was repeated two times and a representative figure is shown.

**Supplementary Table 1.** Carbon sources used in the experiment.

**Supplementary Table 2.** List of chemicals used for HMO and metabolite standard curves.

## Notes

### Competing Interest Statement

The authors have declared no competing interest.

