## Supplementary figures and images for "Time-resolved growth of diverse human-associated Akkermansia on human milk oligosaccharides"

### Supp Figure 1

#### Supp Figure 1

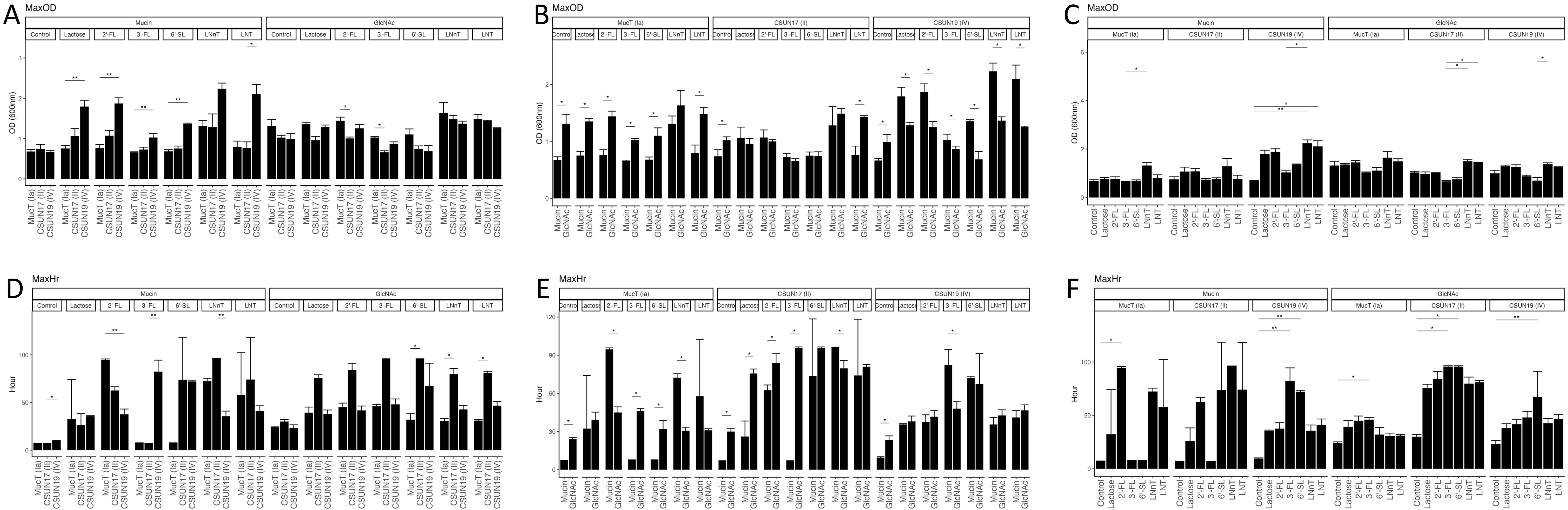

### Supp Figure 2

#### Supp Figure 2

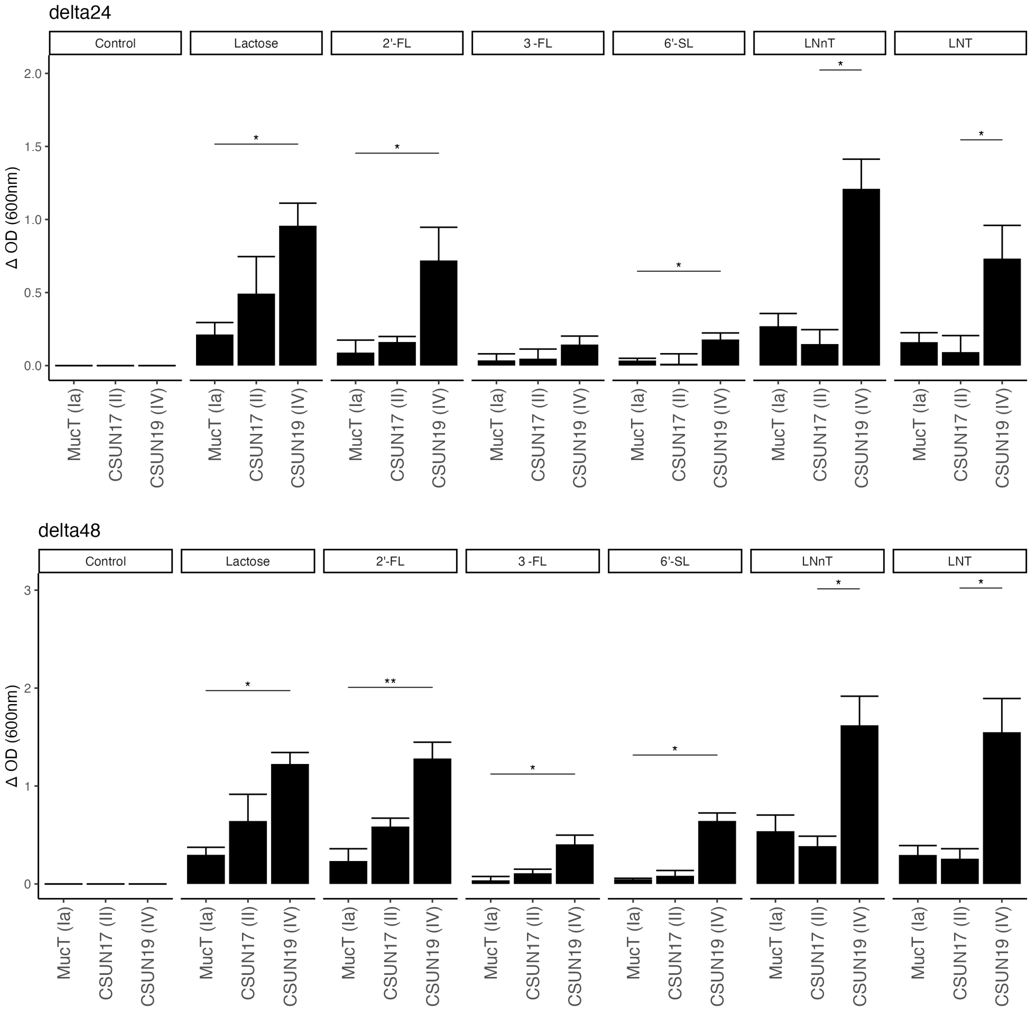

### Supp Figure 3

#### Supp Figure 3

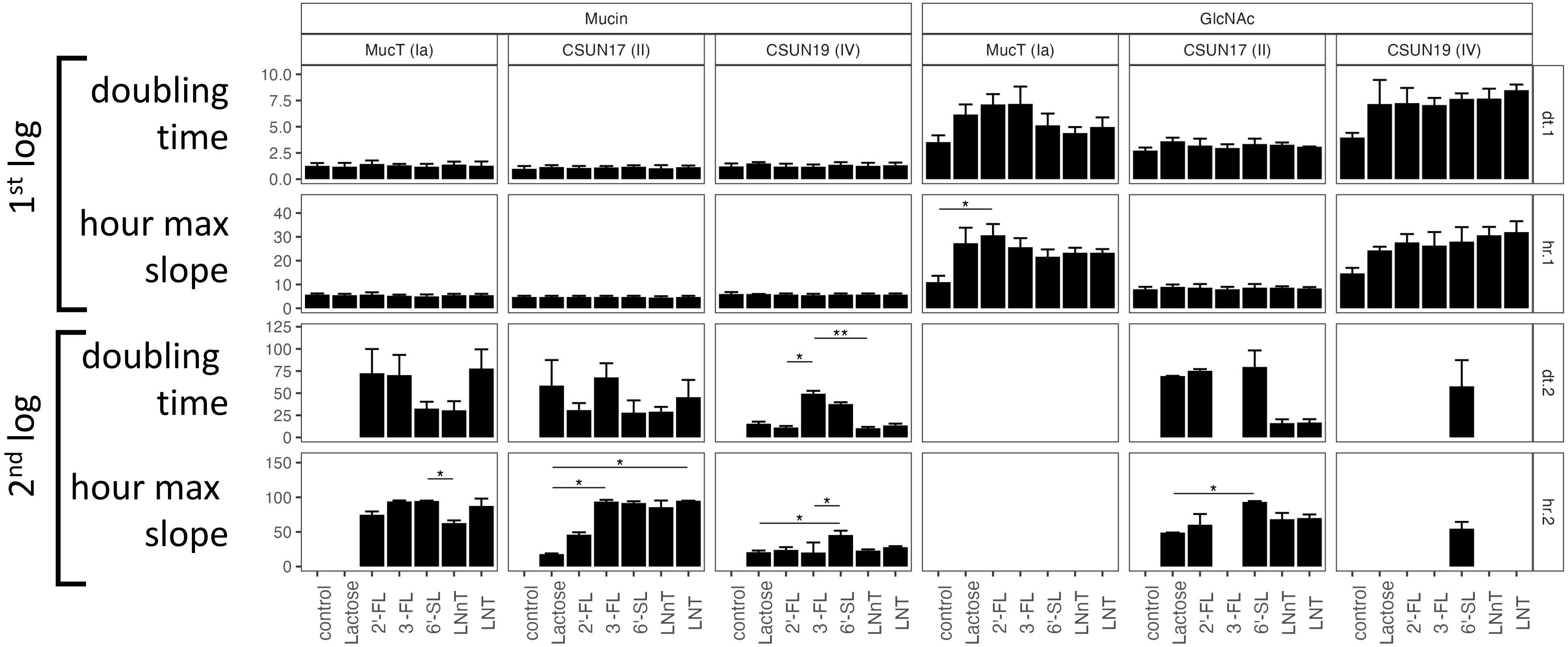

### Supp Figure 4

#### Supp Figure 4

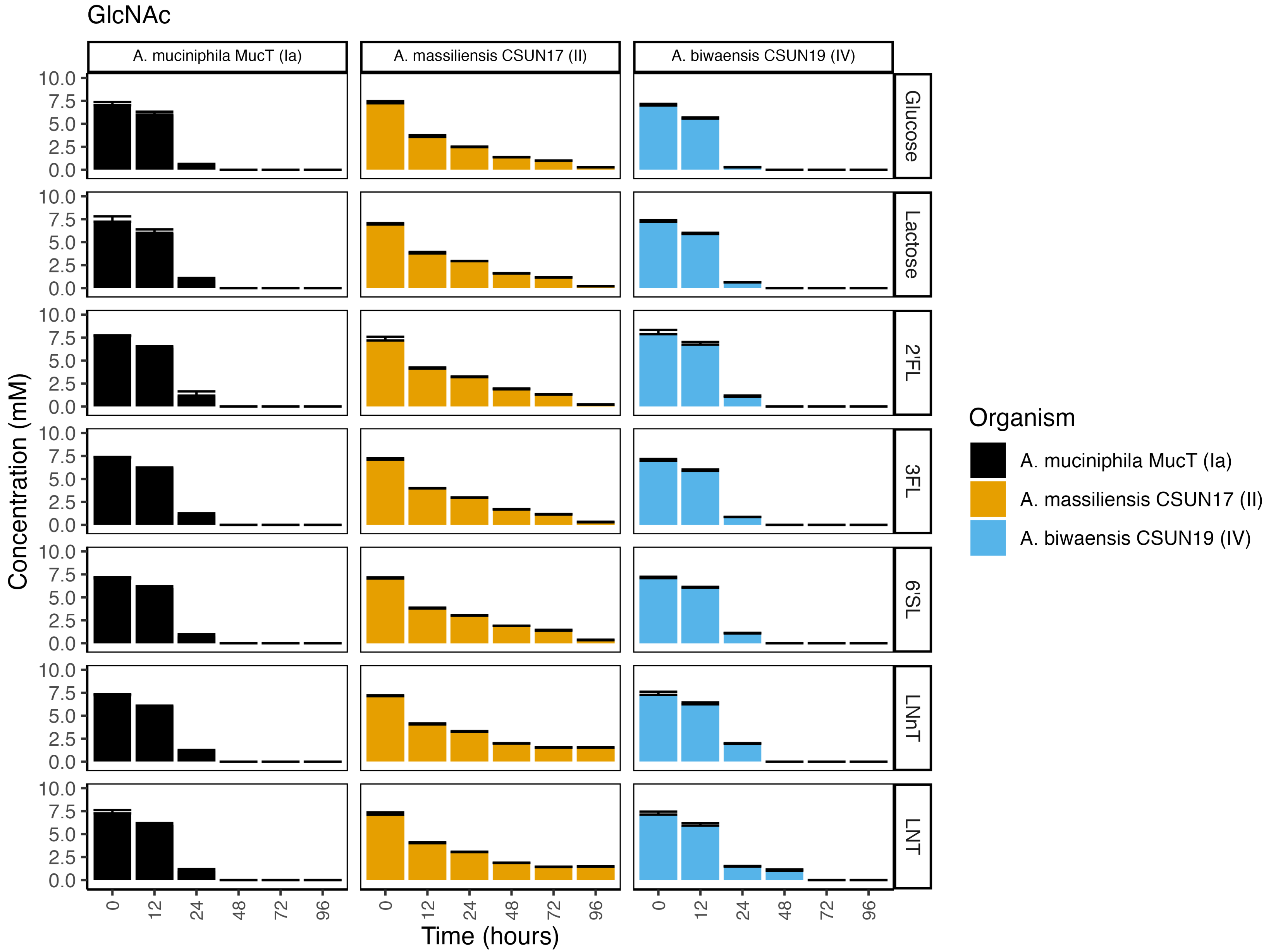

### Supp Figure 5

#### Supp Figure 5

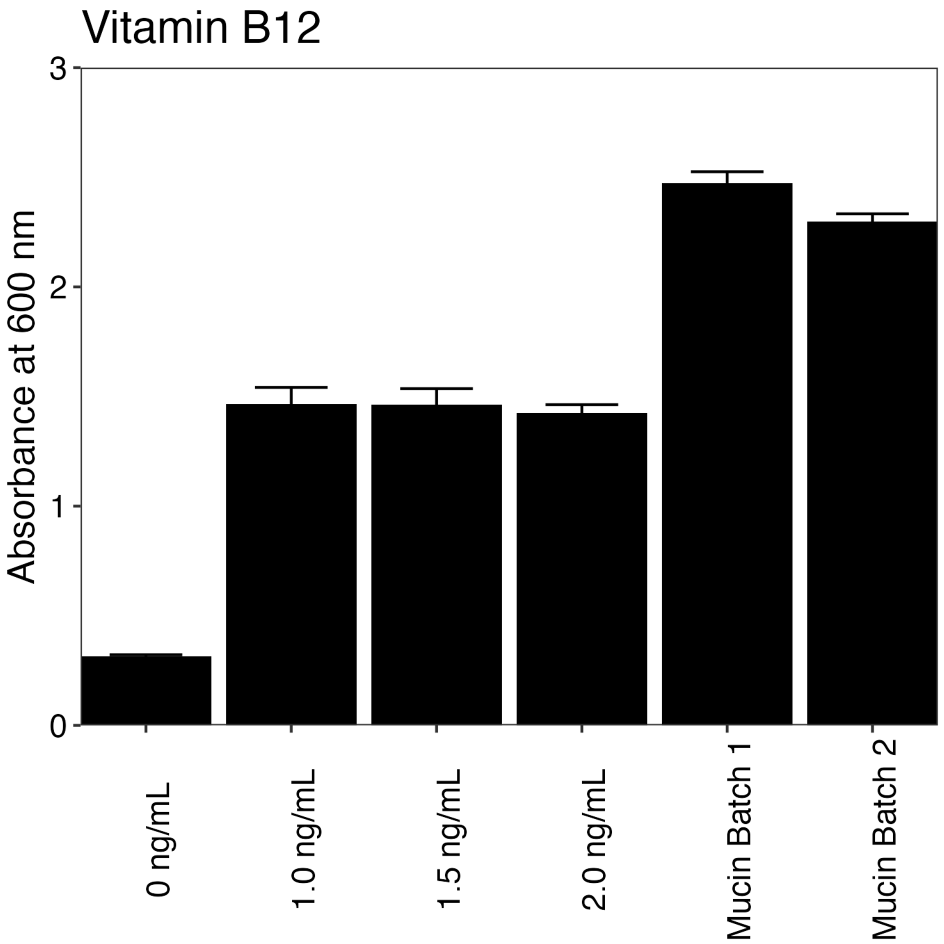
